# Waterlogging shifts ontogenic hormone dynamics in tomato leaves and petioles

**DOI:** 10.1101/2022.12.02.518842

**Authors:** B. Geldhof, O. Novák, B. Van de Poel

## Abstract

Waterlogging leads to hypoxic conditions in the root zone that subsequently cause systemic adaptive responses in the shoot, including leaf epinasty. Waterlogging-induced epinasty in tomato has long been ascribed to the coordinated action of ethylene and auxins. However, other hormonal signals have largely been neglected, despite evidence of their importance in leaf posture control. To adequately cover a large group of growth regulators, we performed a tissue-specific and time-dependent hormonomics analysis. This analysis revealed that multiple hormones are differentially affected throughout a 48 h waterlogging treatment, and, more importantly, that leaf development defines a framework in which this hormonal control is regulated. In addition, we could distinguish early hormonal signals that might contribute to fast responses towards oxygen deprivation from those that potentially sustain the waterlogging response. For example, abscisic acid (ABA) levels peak in petioles within the first 12 h of the treatment, while its metabolites only rise much later, suggesting ABA transport is altered. At the same time, cytokinins (CK) and their derivatives drastically decline during waterlogging in leaves of all ages. This drop in CK possibly releases the inhibition of ethylene and auxin mediated cell elongation to establish epinastic bending. Auxins themselves rise substantially in the petiole of mature leaves, but mostly after 48 h of root hypoxia. Based on our hormone profiling, we propose that ethylene and ABA might act synergistically to dynamically fine-tune the balance of IAA and CK in the petiole, ultimately leading to differential growth and epinasty during waterlogging.

## Introduction

Phytohormones control many developmental processes and are involved in (a)biotic stress resilience, including flooding (Chen et al., 2022). In the model plant *Arabidopsis thaliana*, much progress has been made in elucidating the hormonal framework underlying leaf hyponasty, a hallmark response towards flooding. This upward leaf movement allows plants to reestablish contact between leaves and the air during flooding. Hyponasty also occurs when plants compete with neighbors for light (shade avoidance syndrome; SAS) or escape from radiant heat (thermonasty). The hyponastic response is the result of a complex interplay between several hormones. For example, during shade (low Red/Far-Red), PHYTOCHROME INTERACTING FACTOR 4 (PIF4) and PIF5 signaling induces auxin production, primarily at the leaf tip (Michaud et al., 2017; Pantazopoulou et al., 2017). Auxins are then transported towards the petiole, where they are locally redistributed to the abaxial side. Next, auxin signaling in the lower side of the petiole stimulates local gibberellic acid (GA) biosynthesis, initiating abaxial cell elongation, and hence petiole hyponasty (Küpers et al., 2022). Concomitantly, abscisic acid (ABA) signaling stimulates hyponasty, potentially through the interaction with sugar signaling (Michaud et al., 2022). Much of the SAS-related hormonal crosstalk is shared with thermonastic responses. In this case, temperature sensing through PHYTOCHROME B (PHYB) (Jung et al., 2016; Legris et al., 2016) activates the light signaling cascade through PIFs to modulate thermomorphogenesis, including hyponasty (Park et al., 2019). Intriguingly, the same light signaling pathway is activated in *R. palustris*, undergoing hyponastic growth when flooded (van Veen et al., 2013). In this case, internal accumulation of the gaseous hormone ethylene activates hyponastic bending (van Veen et al., 2013), potentially by inducing auxin redistribution (Cox et al., 2004). In addition, ethylene stimulates differential petiole elongation through reorganization of cortical microtubules (Polko et al., 2012), possibly via brassinosteroid (BR)-dependent signaling, to establish leaf hyponasty (Polko et al., 2013).

Such a hormonal framework has not yet been devised for leaf epinasty or downward leaf bending. Epinasty is a characteristic response to waterlogging or low-oxygen conditions in the roots of species such as tomato. During waterlogging, 1-aminocyclopropane-1-carboxylic acid (ACC), the immediate precursor of ethylene, accumulates in the roots (Olson et al., 1995; Shiu et al., 1998) and is transported upwards to the shoot (English et al., 1995; Jackson & Campbell, 1976), where it is redistributed and converted into ethylene by different members of the ACC-oxidase (ACO) family (English et al., 1995). In the petiole, ethylene likely prompts the redistribution of auxins to induce differential elongation, leading to epinastic bending (Lee et al., 2008). Waterlogging also reduces the levels of GA and cytokinins (CK) in the transpiration stream of tomato, and these hormones seem to counteract the epinastic response (Jackson & Campbell, 1979), but a formal link with other hormones has not yet been established. For example, BRs might be involved in leaf angle control, as the Micro-Tom mutant, defective in BR biosynthesis, has a significantly reduced responsiveness in terms of ethylene-induced epinasty (Edelman & Jones, 2014). Furthermore, the stress hormone ABA is also involved in waterlogging responses. It was shown that ABA levels decline in the xylem during waterlogging (Else et al., 1995), while they rise in the leaves (De Ollas et al., 2021; Hiron & Wright, 1973). Despite these changes, the role of ABA in leaf epinasty has not yet been investigated.

In this paper, we performed a dynamic hormone profiling of tomato leaves and petioles of different ages during waterlogging. We revealed that different hormones are significantly affected during the treatment and that this effect is strongly developmentally regulated. Furthermore, we validated the regulatory role of ABA and CK in the waterlogging-induced epinastic response of tomato.

## Materials and methods

### Plant material and growth conditions

Tomato plants *(Solanum lycopersicum)* of the cultivar Ailsa Craig, M-82 and mutant lines were germinated in soil and afterwards transferred to rockwool blocks. Seeds of the *ait1.1* line, in the M-82 background, (Shohat et al., 2020) were kindly provided by Prof. David Weiss. In addition, the ABA deficient mutant *Notabilis (Not)* was used in the Ailsa Craig background. Tomato plants were grown either in the growth chamber or in the greenhouse until they reached the eighth leaf stage, after which they were used for experiments. The growth chamber climate was set at 18 °C – 21 °C with a constant relative humidity of 65 % and a LED light intensity of 120 μmol s^-1^ m^-2^ (16 h/8 h day/night cycle). In the greenhouse, temperature was set at 18 °C with a relative humidity between 65 – 70 %. Additional illumination was applied if solar light intensity dropped below 250 W m^-2^ (SON-T). Plants received regular fertigation.

### Waterlogging treatment

At the eighth leaf stage, tomato plants were subjected to a waterlogging treatment by flooding the rockwool blocks in individual trays until the oxygen in the root zone was naturally depleted. The treatment was maintained for 24 – 72 h, after which the plants were reoxygenated. Control plants received fertigation throughout the treatment.

### Hormone profiling

Concentrations of the major phytohormones (ABA, IAA, CK, GA, SA, JA) and some of their conjugates (see results section), were quantified in intact leaves and 2 cm petiole segments of leaf 1, 3, 5 and 7 with leaf 1 being the oldest and leaf 7 the youngest (5 biological replicates per treatment). After 12 h, 24 h and 48 h of waterlogging, these tissues were snap frozen, pulverized and analyzed following an UHPLC-ESI-MS/MS protocol described in Šimura et al. (2018).

### BAP treatment

6-Benzylaminopurine (BAP; also named benzyl adenine, BA), a synthetic cytokinin, was dissolved in 1 M KOH and afterwards suspended in lanolin at a final concentration of 1 mg mL^-1^. Lanolin paste was applied to the first 0.5 – 1 cm of the petiole of each leaf.

### Leaf angle measurement

Throughout the waterlogging treatment and during reoxygenation, leaf angle dynamics were monitored in real-time using leaf angle sensors (Geldhof et al., 2021).

### Statistics

All analyses were performed in R (R Core Team, 2019). Interactions between hormones and leaf angles were investigated using principal component analysis (PCA), factor analysis of mixed data (FAMD) and correlation analyses. Effects of treatments and leaf age were determined with one-way or two-way ANOVA and significant differences were either based on a Wilcoxon rank-sum test (α = 0.05) or a Kruskal-Wallis test followed by Dunn’s test (α = 0.05 with and without Bonferroni correction) for single or multiple comparisons respectively. Hormone levels were normalized within a hormone and tissue type to compare relative changes of all hormones between treatments and leaf age classes.

## Results

### Waterlogging induces a tissue-specific and age-dependent hormonal shift

To determine the relative importance of some major plant hormones in regulating waterlogging responses in tomato leaves, we performed UHPLC-ESI-MS/MS on petiole and leaf tissue at three time points (12, 24, and 48 h) and for 4 different leaf ages (leaf 1, 3, 5 and 7). The normalized results of the hormonomics analysis were structured and visualized using PCA (Figure 1). Most of the variation of the hormone profile was related to tissue-specific (PC1) and leaf age-specific (PC2) variability (Figure 1A – B), as validated by factor analysis of mixed data (FAMD) (Supplemental Figure S1). These hormone profiles are however not constant in time, probably due to circadian rhythms (Figure 1B). Within this framework, waterlogging mainly induced a positive and negative shift along PC1 and PC2 respectively (Figure 1A), indicative of a differential hormonal profile depending on ontogeny and tissue-type. These latent axes of variation were associated with certain groups of hormone metabolites (Figure 1C – D): PC1 was mainly associated with ABA and its active catabolites (PA and neoPA), the auxin precursor TRP and certain CK catabolites, including cZ7G and iP7G (irreversible catabolites) and tZOG and tZROG (reversible catabolites; but opposite effect), and active JA and JA-Ile; PC2 mainly represented GA19 and IAA and some of its precursors (ANT and TRA). Hormones associated with both PCs potentially contribute to aforementioned waterlogging-induced changes, and this shift seems to largely govern active CKs and their riboside precursor (cZR, tZR and iPR) and conjugates (cZ7G, iP7G, tZOG, DHZ7G).

**Figure 1:**
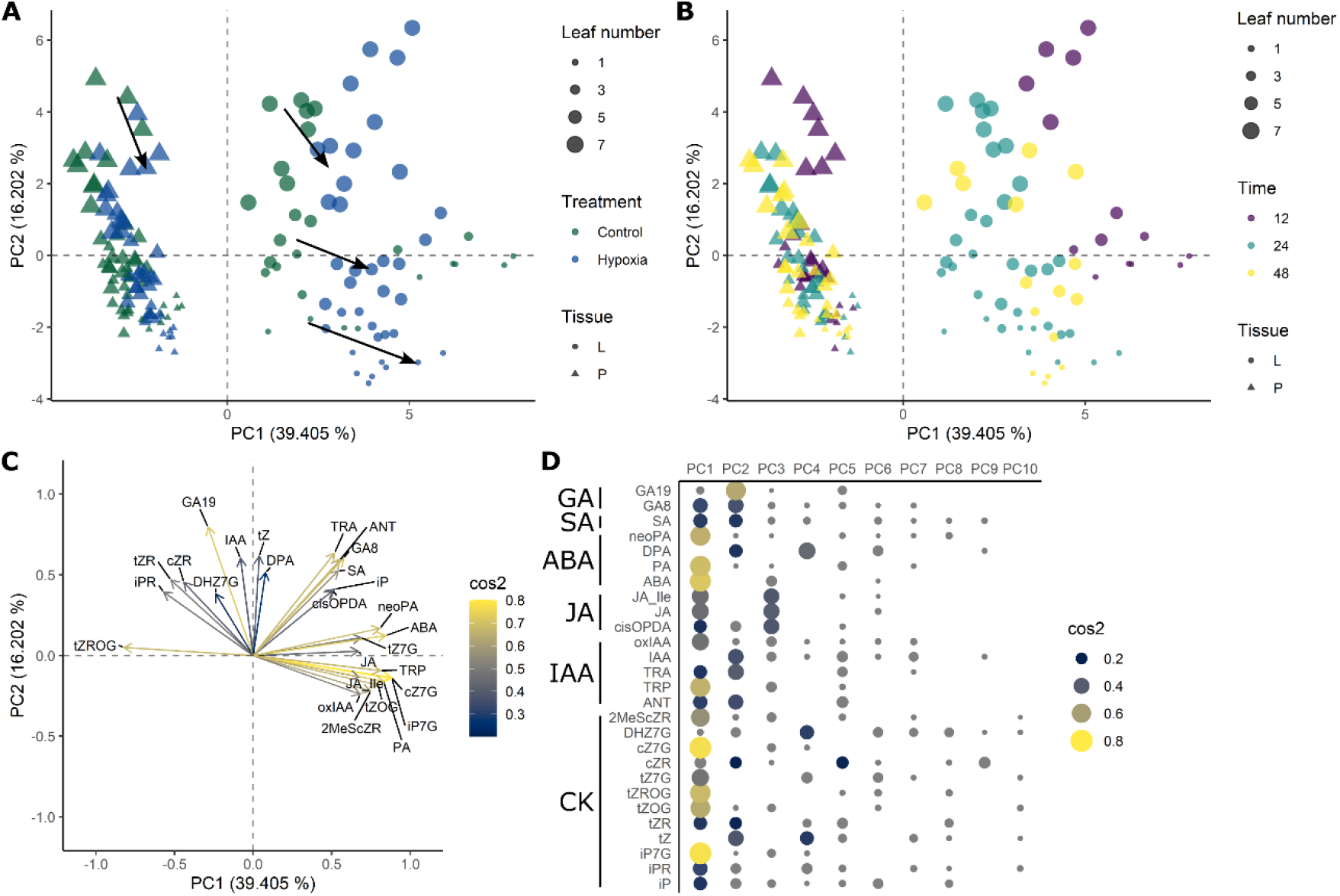
Hormonal changes in tomato leaves and petioles during waterlogging. PCA performed on the hormone levels in leaf (L) and petiole (P) tissue of different ages (leaf number 1, 3, 5 and 7) in relation to (A) the waterlogging treatment and (B) the sampling time (12, 24 and 48 h). Arrows in (A) indicate the waterlogging effect. (C) PCA variable plot depicting possible interactions between hormones and their representation within PC1 and PC2. (D) Quality of representation (cos2) of the different variables in PC1 – PC10.

Next, we determined the waterlogging effect on individual hormone levels, normalized per metabolite and tissue type (Figure 2A). Treatment effects largely corroborated the overall trends from the PCA, but also provided some additional insights. ABA derivatization to PA and consecutively DPA seem to be enhanced in mature and young leaves respectively (Figure 2A), suggesting fast conversion of PA to the inactive DPA in the latter. ABA levels themselves increased rapidly in petiole tissue within 12 h of the waterlogging treatment. On the other hand, JA levels decreased in leaves of all ages at this early time point, while SA levels only dropped in old leaves (leaf 1) after 12 h of waterlogging. IAA levels and those of its precursor TRP rise mainly later, after 48 h of waterlogging, and especially in petioles of mature leaves. In contrast, levels of active CK (tZ and iP) and their riboside precursors (tZR and iPR) drop in leaves and petioles in an age-dependent way, being more pronounced in young leaves. As we were unable to detect active GA’s in our analysis, no general conclusions can be drawn on the effect of waterlogging on leaf and petiole GA homeostasis.

**Figure 2:**
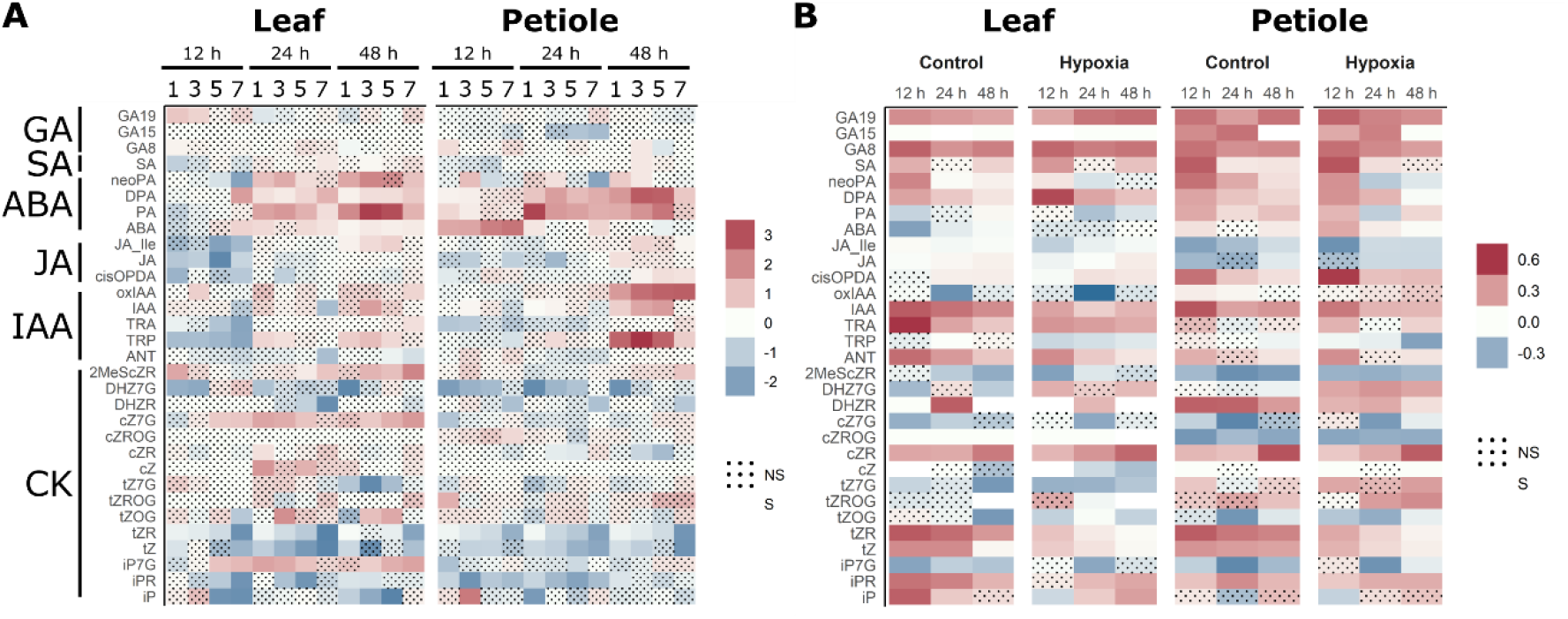
Ontogenic hormone profiles in control condition and during waterlogging (12, 24 and 48 h) in tomato petioles and leaves of different ages (leaf 1, 3, 5, 7). Effect of (A) waterlogging and (B) leaf age on normalized hormone levels. A positive effect in (A) indicates an increase in hormone levels during waterlogging and a positive effect in (B) indicates higher hormone levels in younger leaves Significant effects are indicated as non-marked tiles and were determined with ANOVA (Kruskal-Wallis α = 0.05).

### Development sets the scene for hormone dynamics during waterlogging

Our PCA analysis revealed a strong link between the hormonome and leaf age, indicating that hormone homeostasis is dynamically regulated throughout development (Figure 1). This ontogenic association prompted us to quantify the actual leaf age effect on individual hormone levels (Figure 2B). In control conditions, young leaves have a relatively higher content of GA19, GA8, IAA (and its precursors TRA and ANT) and active CKs (tZ and iP) and CK ribosides (cZR, tZR and iPR; see also Figure 1). Some CK conjugates, including the N-glucosides cZ7G, tZ7G and iP7G are generally less abundant in younger leaves in control conditions. Although hormone profiles of the petiole mostly overlap with those of leaves, some hormones seem to change depending on the tissue-type. For example, JA and JA-Ile levels do not change in leaves, but increase with age in the petiole, while the JA precursor cisOPDA decreases with age in the petiole. The opposite is true for ABA, which increases in aging leaves, but decreases in aging petioles (mostly at 12 h). This tissue-specific ontogenic regulation is also dynamically regulated through the day (12 h vs 24 h in control conditions), pointing towards compartmentalization of these processes in both space and time, possibly through transport or circadian effects.

If developmental differentiation orchestrates hormone homeostasis, ontogeny might as well regulate hormonal shifts during waterlogging. Indeed, already after 12 h of waterlogging, leaf age significantly affects hormone dynamics in both leaves and petioles. Upregulation of ABA levels by waterlogging was initially (12 h) confined to petioles and young leaves, and later (24 and 48 h) followed by an upregulation in leaves of all ages (Figure 2A). Intriguingly, the formation of PA (an active conjugate of ABA) is dampened in young leaves (leaf 7) as compared to older leaves, while its conversion into DPA remains relatively high. This illustrates a complex interplay between ABA synthesis and catabolism in regulating tissue-specific, age- and time-dependent waterlogging responses. The IAA content was reversely affected in leaves of different ages: IAA levels significantly dropped in young leaves (leaf 7) after 24 h of waterlogging treatment, while they increased significantly in older leaves (48 h) and petioles (12 – 48 h). Together with the fast (12 h) reduction of its precursor TRP mainly in old and mature leaves during waterlogging, this suggests a transition from IAA synthesis towards an elevated supply of auxins, possibly mediated by IAA redistribution within the plant, leading to a peak of TRP and IAA in the petiole at 48 h of waterlogging.

### CK metabolism as a target for ontogenic and waterlogging-induced changes

Although auxins and CKs are often assumed to act antagonistically, biosynthesis of the main CKs and their derivatives are subject to multiple layers of regulation, leading to complex homeostasis (Figure 3). Based on our ontogenic profiling, it has become clear that certain CKs are downregulated while others are not, and that this regulation is tissue-specific (Figure 2A and Figure 3A). The active CKs iP and tZ were more abundant in leaves or both leaves and petioles respectively and decreased with leaf age. While the reduction of tZ during waterlogging was clearly more pronounced in young tissue, this shift was more complex for iP. The CK ribosides cZR, tZR, iPR (and DHZR) were predominately detected in petioles, and declined both with leaf age and relatively more in young leaves during waterlogging. The *N*-glucosides cZ7G, tZ7G were elevated in mature and older leaves and either increased in all or decreased in old leaves during waterlogging. iP7G also increased with leaf age, and increased in leaves throughout the treatment.

**Figure 3:**
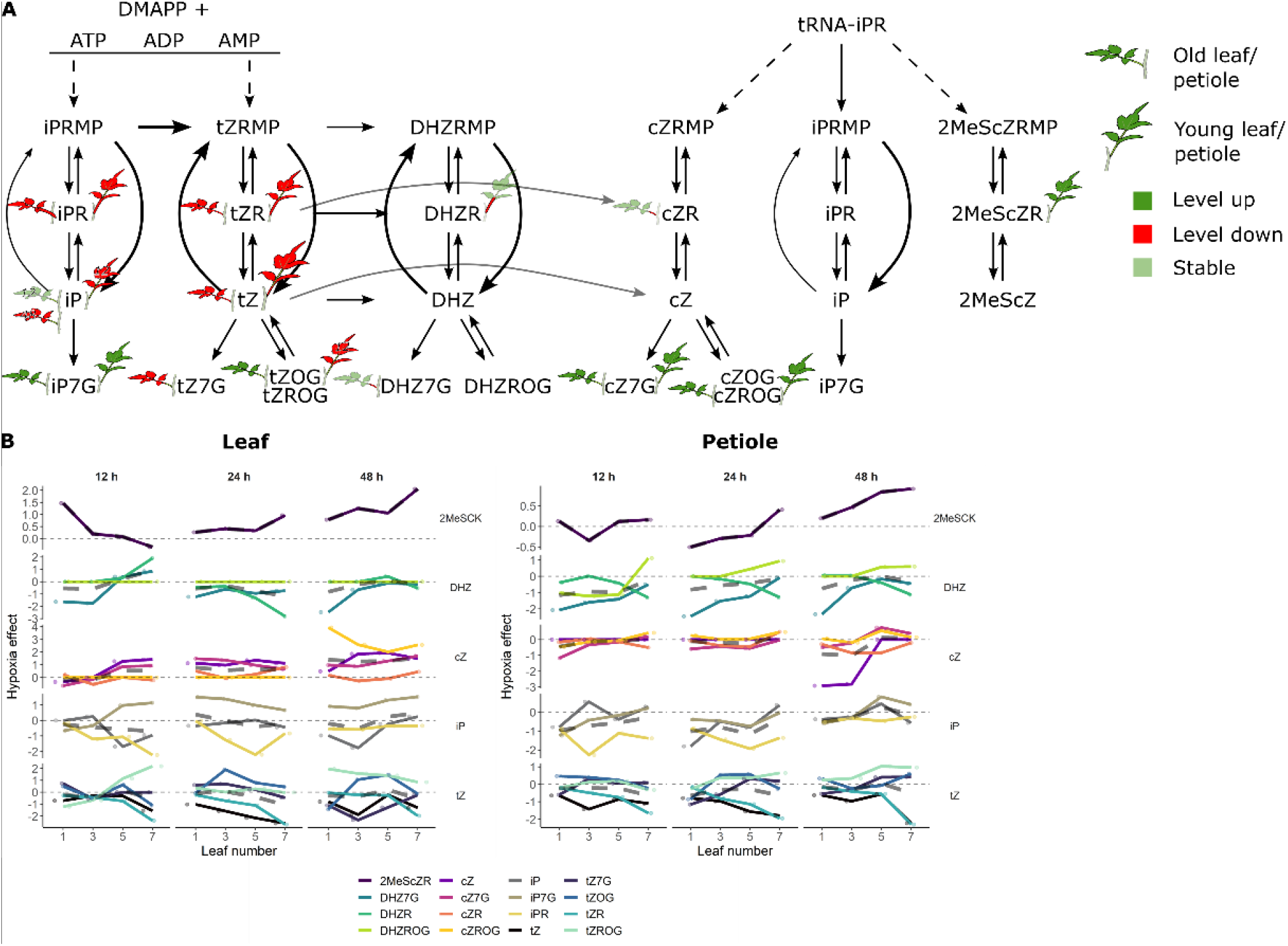
Ontogenic CK dynamics during waterlogging. (A) CK biosynthesis pathway, including major CK precursors or conjugates detected in this study, represented with a leaf color scale. Changes are indicated for old/mature and young leaves for both the leaf and petiole. The size of the petiole indicates basal levels (especially for ribosides). (B) Effect of waterlogging treatment on ontogenic CK trend in leaves and petioles. A positive effect indicates increase of the respective CK type.

Ontogenic divergence of CK levels during waterlogging defines a possible mechanism orchestrating developmental plasticity (Figure 3B). In general, cZ-type CKs increase faster in young leaves after the start of the waterlogging treatment, followed by a slower rise in older leaves. DHZ CKs (DHZ7G) mainly seem to be reduced in older leaves, while iP- and tZ-type CKs increase and decrease depending on the specific type and the developmental stage. Interestingly, transportable CK types, including the ribosides iPR, tZR, DHZR and cZR are consistently downregulated mainly in petiole tissue, which could suggest that both local production and transport of most CKs is inhibited. These CK dynamics largely coincide with the results from our PCA analysis (Figure 1).

### Hormone homeostasis associates with angular dynamics during waterlogging

Changing hormone levels might not directly evoke morphological changes in the leaf petiole during waterlogging. To explore the relationship between hormone profiles and leaf angle changes, we investigated the age-specific effect of waterlogging on epinastic bending. Using a real-time angle sensor (Geldhof et al., 2021), we quantified the epinastic movement during waterlogging and subsequent reoxygenation (Figure 4A). The first changes in angle dynamics occurred after approximately 6 – 8 h of waterlogging, and the leaves reached their maximal angle after 48 – 72 h of waterlogging, short before the end of the treatment. Young leaves (leaf 7) were able to reposition during reoxygenation, while the epinastic bending persisted in mature (leaf 5) and old leaves (leaf 3). Next, we determined correlations between the epinastic leaf angle and the petiole hormone profiles (Figure 4B). This effect seemed not yet significant after 12 h, despite visible changes in the hormonome and leaf angle dynamics (Figure 4A), indicating that petiole bending lagged behind on the actual hormonal changes. As a result, the correlation between instant angular change (12 h) and levels of multiple hormones in the petiole might thus only cover fast responses, while the induction of bending might have occurred at an earlier time (Figure 4C). Notably, hormones that show the strongest changes, including IAA, ABA and CK conjugates, seemed to be associated with the angle change, for these instant responses. ABA itself, for example, was negatively correlated with leaf angle change through the entire treatment (12 – 48 h; total in Figure 4C), despite it showing a fast increase during waterlogging. To gain more insight into these sometimes time-conflicting results, we quantified the effect of individual normalized hormone levels on average leaf angle change (Figure 4C), taking into account the time lag between activator and response (Figure 4C). Even though most of the interactions are not significant, major shifts in the ABA/PA and IAA metabolism and conjugation of CK in the petiole seem to relate to leaf angle changes at a later time point. Interestingly, a decrease of the JA and TRP pool also seem to associate with epinasty. Also SA levels were correlated with the total angular change, probably independently of the treatment (Figure 2A).

**Figure 4:**
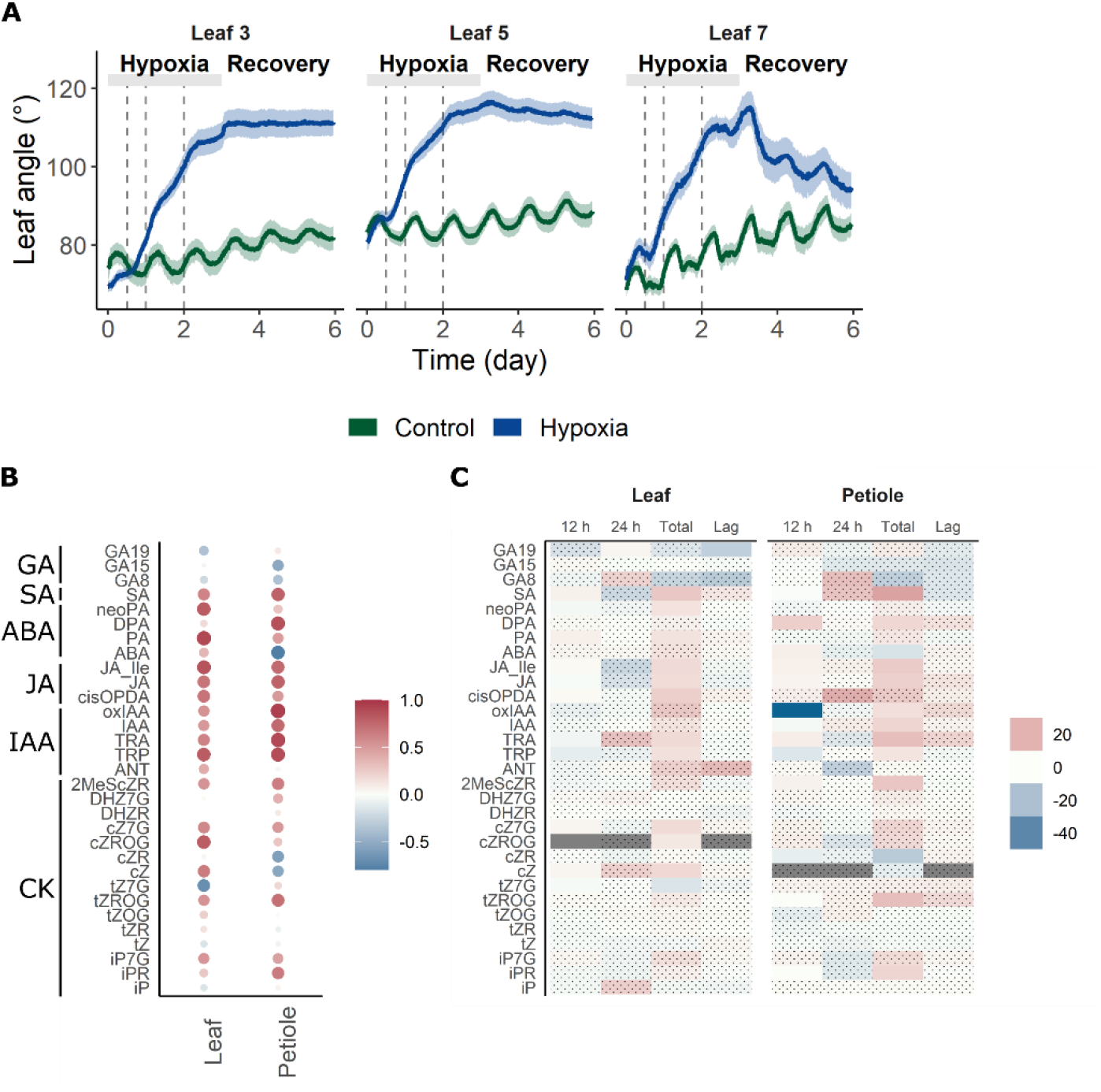
Association between hormone levels and leaf angle changes during waterlogging. (A) Leaf angle dynamics in leaves of different ages during waterlogging and subsequent reoxygenation (n = 10). Dashed lines indicate sampling points for the hormonomics analysis (n = 5). (B) Correlation plot of the different hormone levels in the leaf and petiole and the resulting leaf angle at all time points (12 – 24 – 48 h). Size and color indicate the strength of the correlation. (C) Direct and lagged effect of hormone levels on leaf angle change after 12 h and 24 h of waterlogging or for the entire period (total) for both the leaf and the petiole. Hormone levels were normalized per hormone and tissue type and effects were determined using one-way ANOVA. Shaded effects were not significant in the respective test (α = 0.05).

### ABA modulates the epinastic response in tomato

The relatively fast increase in ABA content and its catabolites PA and DPA at the onset of waterlogging (Figure 5A) prompted us to investigate in more detail the role of ABA in directing leaf epinasty. Therefore, we quantified the leaf angle change of the ABA biosynthesis mutant *notabilis (not)*, defective in the *9-CIS EPOXYCAROTENOID DIOXYGENASE1 (NCED1)* gene, and in an ABA transporter mutant of ABA-IMPORTING TRANSPORTER1 *(ait1.1)*, which has been reported to be less sensitive to the hormone (Kuromori et al., 2018; Shohat et al., 2020).

**Figure 5:**
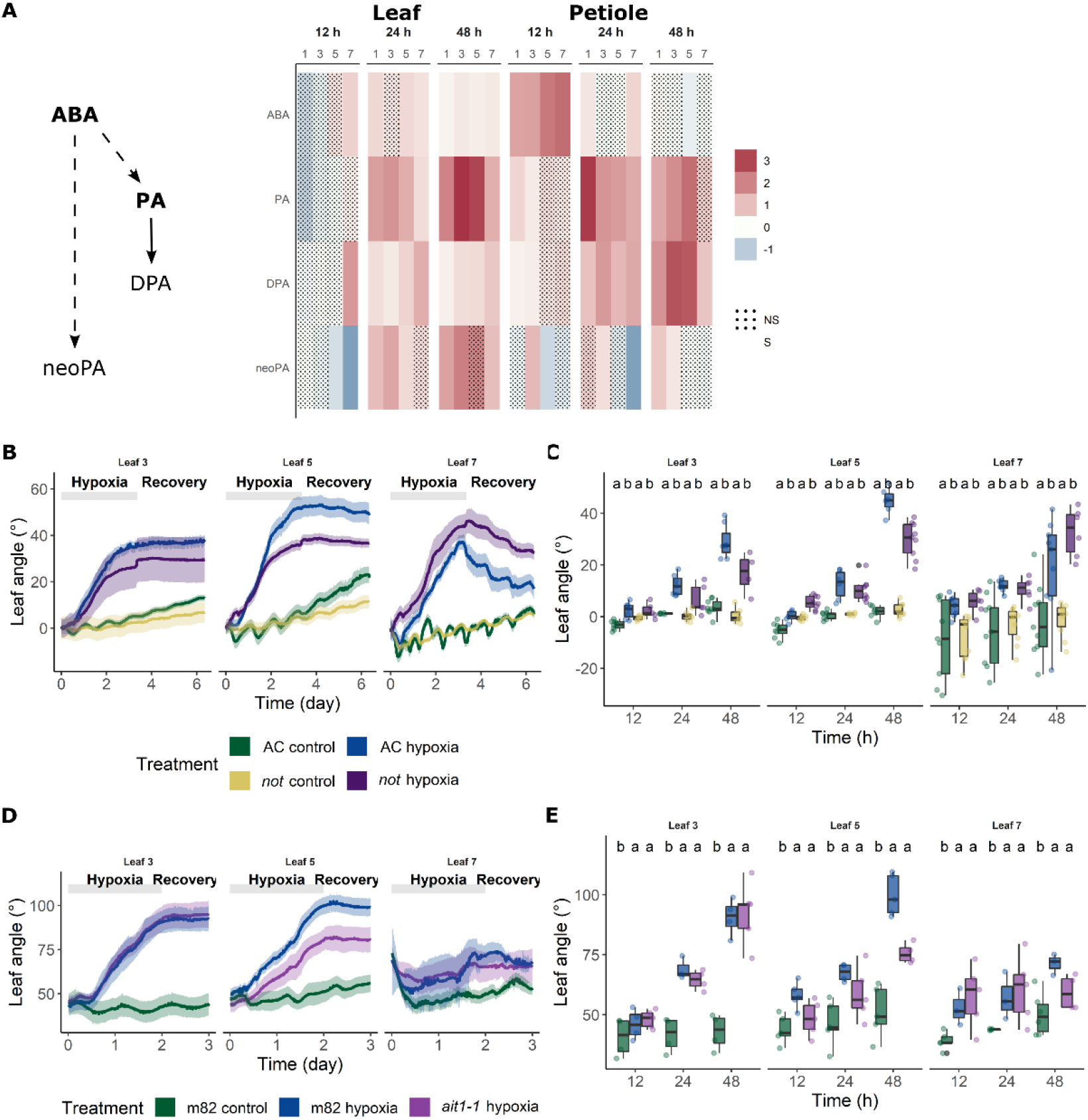
Effect of ABA biosynthesis and transport defects on leaf angle change during waterlogging. (A) Effect of waterlogging on normalized levels of ABA and its catabolites PA, DPA and neoPA. (B – E) Leaf angle change in (B – C) Ailsa Craig and *notabilis* (n = 3 – 10) and (D – E) m82 and *ait1-1* during waterlogging and subsequent recovery (n = 3 – 6). The line and shaded zones in (B & D) represent the average and the 90 % confidence interval respectively. Values in (C & E) represent the average leaf angles at 12, 24 and 48 h of waterlogging. Different letters in (C & E) represent significantly different groups (Dunn’s test, α = 0.05 with Bonferroni correction).

The *not* mutation had an age-dependent impact on leaf posture during waterlogging (relative to the angle at the start of the treatment; Figure 5A – B), with a seemingly reduced and enhanced epinastic bending in mature (leaf 5) and young leaves (leaf 7) respectively. These young leaves also showed slower repositioning during reoxygenation. The rather moderate effects suggest that ABA primarily modulates the epinastic response, or that the *not* mutant still retains some ABA responsiveness during waterlogging. However, the reduced ABA import capacity of the *ait1-1* mutant leads to a similar trend in mature leaves (significant without Bonferroni correction), indicating that disrupted ABA transport might dampen the epinastic response (Figure 5C – D).

### Local supply of CK at the petiole effectively abolishes leaf epinasty

In contrast to ABA, waterlogging strongly reduces active CK levels and their transportable precursors in both leaves and petioles (Figure 6A – B; tZ and iP), indicating CK might be involved in regulating leaf posture. To investigate the role of CK in leaf bending, we applied lanolin paste containing benzylaminopurine (BAP, a synthetic cytokinin) directly to the petiole before the start of the waterlogging treatment and subsequently monitored leaf angle dynamics (Figure 6C – D). The BAP treatment effectively inhibited leaf epinasty in mature (leaf 4; significant in Figure 6D without Bonferroni correction) and young leaves (leaf 5), and enhanced recovery after reoxygenation. This was less the case for older leaves (leaf 3).

**Figure 6:**
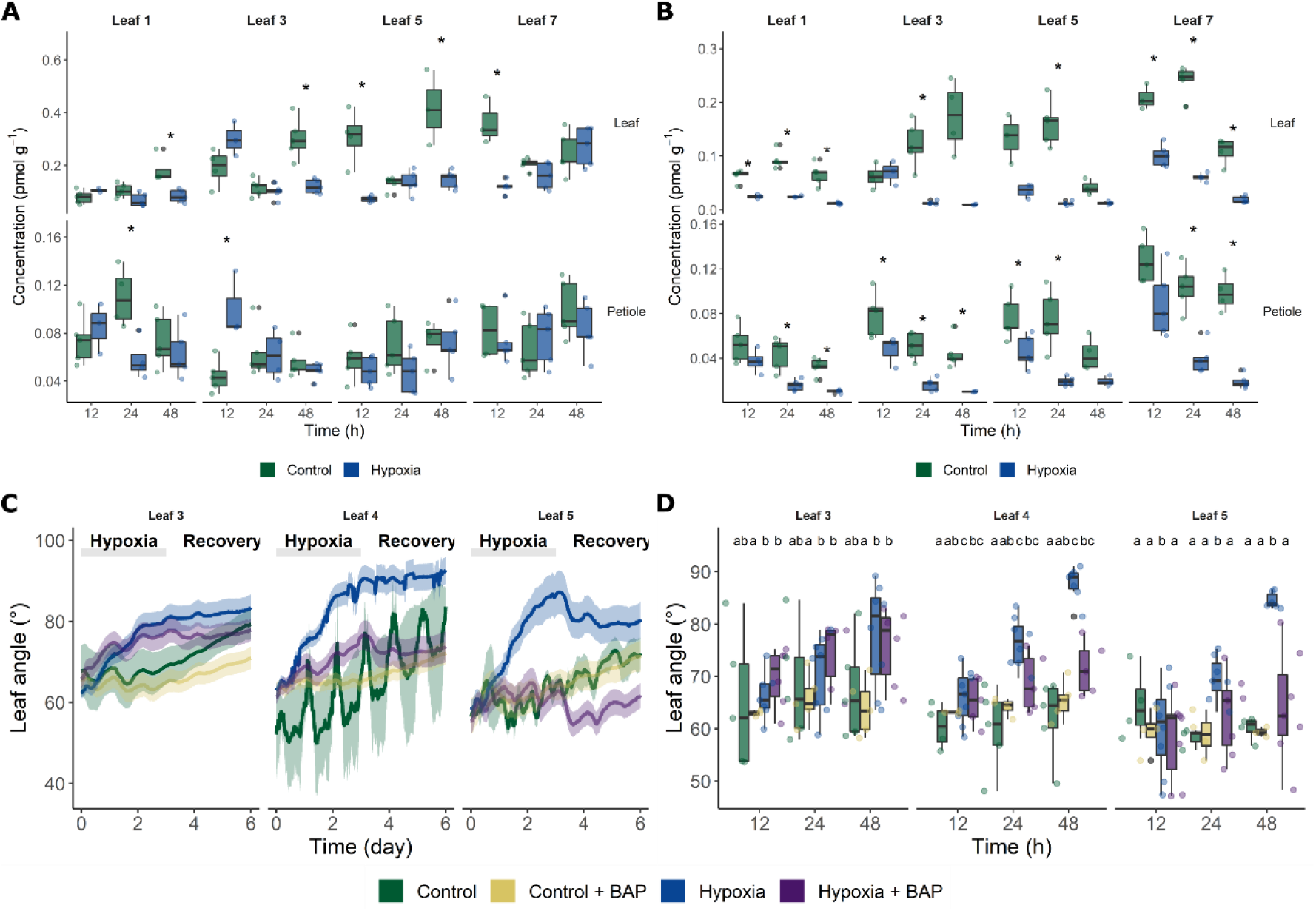
Local application of CK at the petiole reduces epinastic bending during waterlogging. (A – B) Levels of (A) iP and (B) tZ in tomato leaves and petioles during waterlogging (n = 5). (C) Leaf angle dynamics of Ailsa Craig leaves (leaf 3, 4 & 5) with and without BAP treated petioles in control and waterlogging conditions and after reoxygenation (n = 5 – 8). (D) Average leaf angles at 12, 24 and 48 h of waterlogging, with and without BAP treatment. Different letters in (C & E) represent significantly different groups (Dunn’s test, α = 0.05 with Bonferroni correction).

After reoxygenation, young leaves (leaf 5) of waterlogged and CK-treated plants reposition to a more erect state, indicating CK could have an effect on upward movement when returning to normoxic conditions. The application of additional CK activates upwards bending mostly in young leaves (leaf 5), and mildly in older leaves (leaf 4 and to a lesser extent leaf 3), that normally do not recover after waterlogging. Collectively, these results suggest that a local reduction of CK levels in the petiole is needed to initiate leaf bending during waterlogging and that their increase during reoxygenation could restore leaf position. In addition, this CK homeostasis seems to be ontogenically regulated.

## Discussion

### Beyond ethylene: fast responses to waterlogging include shifts in ABA and JA levels

It is well established that waterlogging induces the accumulation (Olson et al., 1995; Shiu et al., 1998) and transport (English et al., 1995; Jackson & Campbell, 1976) of ACC from hypoxic roots to the shoot, followed by subsequent upregulation of ethylene production in the shoot. Many other hormones have been proposed to play a role in signaling of waterlogging or flooding stress, but their concerted action remains largely elusive (Jackson, 2002). For example, hormones such as ABA, GA, IAA and CK can also be transported via the vascular tissue, allowing them to act systemically, while their local biosynthesis is equally important for adequate stress responsiveness (reviewed in Kuromori et al. (2018) & Li et al. (2021)). We have now shown that waterlogging shifts the hormonome of tomato leaves and petioles, and that this response is highly tissue-specific. This analysis also revealed that both JA and ABA are among the early response hormones during waterlogging stress.

Previously, it was shown that reduced oxygen availability in the rooting zone represses ABA biosynthesis genes and ABA levels in Arabidopsis (Hsu et al., 2011) and tomato roots (De Ollas et al., 2021), leading to a reduced ABA export to the shoot by the xylem (Else et al., 1995). Despite reports about the repression of ABA production by ethylene in species such as *Rumex palustris* (Benschop et al., 2005), local ABA biosynthesis and ABA levels are upregulated in tomato leaves by waterlogging (De Ollas et al., 2021; Hsu et al., 2011), indicating ethylene-ABA crosstalk might be tissue-specific. Indeed, we observed that ABA content increased mostly in the petiole within 12 h of the start of a waterlogging treatment (Figure 2). This suggests that local ABA production or ABA influx contribute to the increasing ABA levels in petioles during waterlogging. In contrast, ABA catabolism to PA and consequently DPA increased only after 24 – 48 h of waterlogging in both the leaf and petiole. In *R. palustris*, the conversion to PA is believed to be part of the ethylene-mediated pathway inducing petiole elongation during flooding (Benschop et al., 2005). Interestingly, the role of PA might actually be more important than anticipated, as it interacts with at least a subset of ABA receptors (Weng et al., 2016).

Besides ethylene-ABA cross-talk, ABA-signaling also influences JA levels. Recently, it was shown that JA levels decline in tomato leaves during 24 h of waterlogging, potentially as a result of ABA signaling (De Ollas et al., 2021). Our results corroborate these findings and reveal that this JA drop in leaves is transient and only significant within the first 12 h of waterlogging, similar to the ABA spike in the petiole.

### Are ABA and its transport involved in waterlogging-induced differential elongation?

The rapid increase (12 h) of ABA levels mainly in petiole tissue during waterlogging raises the question where this ABA originates from. While increased import through the xylem from the hypoxic root system is unlikely (Else et al., 1995), there are some indications for ontogenic control of ABA flows. After 12 h of waterlogging, ABA levels in older leaves decreased (not significantly), while the youngest showed an increase in ABA levels, suggesting a potential ABA transfer from source to sink leaves respectively (Figure 7). This type of ABA recirculation has been described in castor bean *(Ricinus communis)* during salinity stress (Jeschke et al., 1997) and in *Xanthium strumarium* during drought stress (Zeevaart & Boyer, 1984).

**Figure 7:**
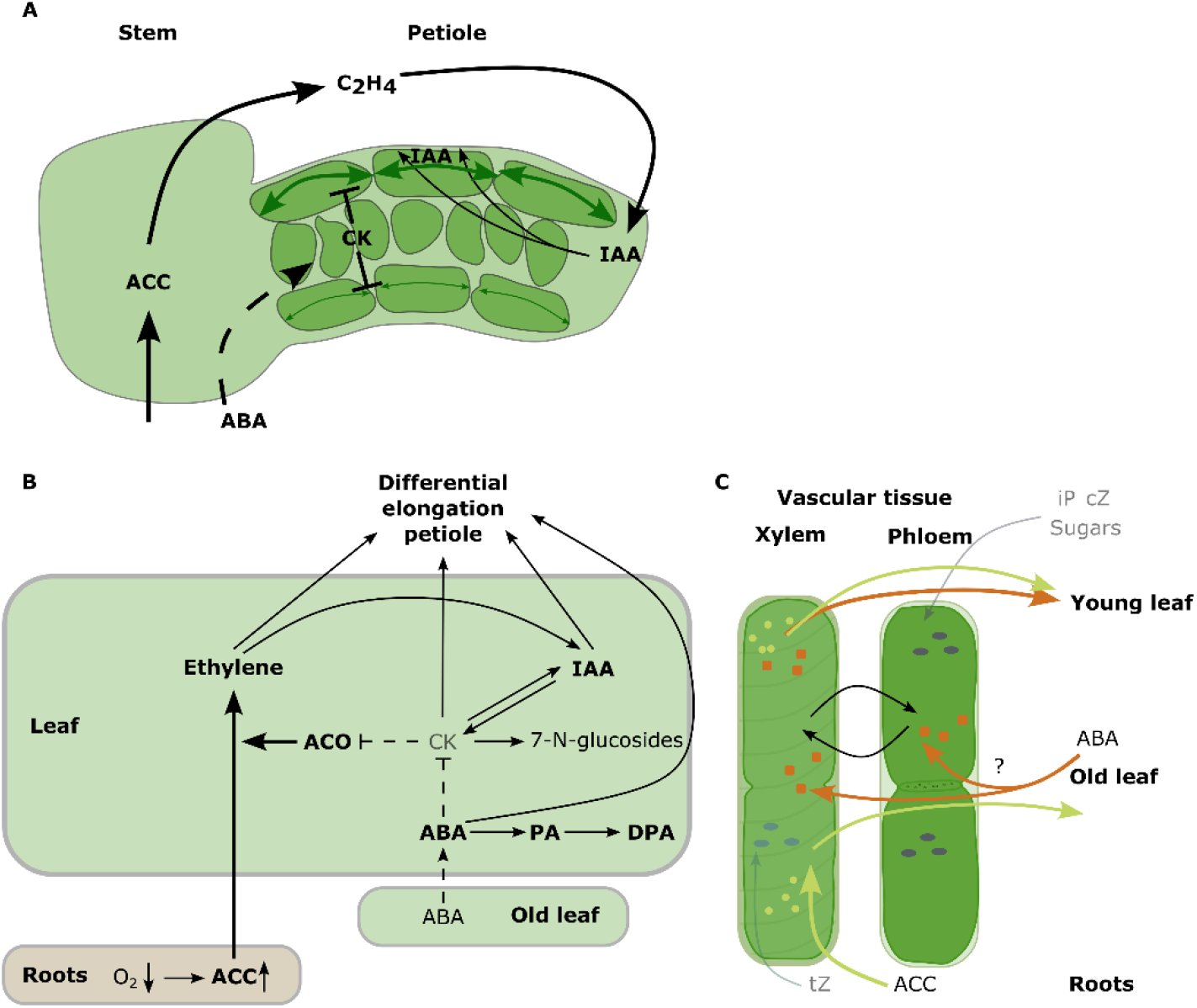
Schematic overview of the systemic hormonal changes in tomato (A) petioles, (B) leaves and (C) vascular tissue during waterlogging. ACC accumulation and its transport from the roots is greatly enhanced, leading to a large influx in the leaves, where it gets converted into ethylene. At the same time, ABA is translocated through the plant, likely from older source leaves to the other leaves. In contrast, both active CK pools and CK transport are reduced and their *N*-glucosylation stimulated. The rapid increase in ethylene production, together with fast ABA accumulation and a CK reduction in the petiole, releases a break mechanism, allowing for auxin accumulation (probably due to a reduced IAA export from the leaf) that regulates differential elongation and asymmetric growth leading to the downwards bending of the petiole.

Although we cannot rule out increasing ABA levels by local biosynthesis, both systemic and local ABA transport can contribute to epinastic bending during waterlogging. The endogenous ABA levels are the result of the coordinated action of ABA exporters and importers, one of which is NITRATE TRANSPORTER1/PEPTIDE TRANSPORTER FAMILY4.6 (NPF4.6), also known as ABA-IMPORTING TRANSPORTER (AIT1) (reviewed in Kuromori et al. (2018)). AIT1 localizes to guard cells (Shimizu et al., 2021), but also to the vascular bundles (Kanno et al., 2012; Shohat et al., 2020), alongside major ABA biosynthesis enzymes (Kuromori et al., 2014). The presumably reduced ABA import capacity in the *ait1.1* mutant might alter ABA translocation, and thus dampen the ABA responses. This might in turn explain the reduction in epinastic bending in the *ait1.1* mutant during waterlogging (Figure 5).

ABA is often described to counteract ethylene biosynthesis and action. Ethylene enhances elongation during flooding by inactivating the ABA metabolism in *R. palustris* (Benschop et al., 2007) and rice *(Oryza sativa)* (Saika et al., 2007), indicating that ethylene and ABA might act antagonistically in regulating elongation (Zeevaart, 1983). In tomato, mutations in ABA biosynthesis release suppression of ethylene mediated processes, which in turn leads to ethylene-associated growth (Sharp et al., 2000). Indeed, the *not* mutant in tomato has an enhanced ethylene production and a seemingly epinastic appearance (Thompson et al., 2004). Recent work has provided evidence that the *not* mutation, although yielding a lower basal ABA production, does not seem to confer shifts in shoot ABA biosynthesis during waterlogging (De Ollas et al., 2021) and salinity (Holsteens et al., 2022). However, De Ollas et al. (2021) did not take into account leaf ontogeny, thus neglecting possible effects of ABA translocation. Together with its residual production in the shoot, ABA might be able to locally sustain partial responses such as differential elongation and epinasty (Figure 7).

While ABA overproducing tomato mutants are less epinastic, they also have longer petioles (Thompson et al., 2007), indicating ABA could indeed stimulate growth in the petiole. Moreover, exogenous ABA applications have a stimulatory effect on epinastic bending in lemonwood *(Pittosporum eugenioides)* (Dwyer et al., 1995), indicating ABA can also activate differential growth. Altogether, our assay demonstrates that ABA could play a role as an early stress signal directly interfering with morphological plasticity in an age-dependent way.

### Petiolar auxin levels peak during late waterlogging responses

Our hormonomics analysis revealed that auxin levels are also upregulated during waterlogging, but mostly at later time points (24 – 48 h). The initial (12 h) decrease of IAA precursors (TRP and TRA) in leaves did not coincide with a rise in IAA levels, indicating TRP might have been repurposed for either protein synthesis or specialized metabolism, such as melatonin and serotonin production (reviewed in Bhowal et al. (2021); Hardeland (2016)). Although an increase of TRP has been reported in barrel clover *(Medicago trunculata)* during waterlogging (Lothier et al., 2020), little is known about its direct effects on the actual stress response. TRP might be involved in leaf movements, as it was shown that TRP can regulate pulvinar motor cell functioning in *Mimosa pudica* (Dédaldéchamp et al., 2019). In addition, TRP has been shown to directly interfere with plant and cell growth in Arabidopsis (Jing et al., 2009).

The intricate coordination of auxin with cytokinin metabolism, transport and signaling is one of the core axes regulating normal plant growth and development (reviewed in Schaller et al. (2015)). This interaction between both hormones seems to be largely determined by the developmental stage (Nordström et al., 2004). Indeed, young tomato leaves harbor the largest pool of both auxins and active CK (Figure 2), suggesting ontogenic control of their action. The potential of auxins and CK to co-regulate developmental processes is exemplified by SYNERGYSTIC ON AUXIN AND CYTOKININ1 (SYAC1), which is at the crossroads of auxin and CK signaling (Hurný et al., 2020). SYAC1 acts through tissue- and age-dependent mechanisms to orchestrate processes such as primary root growth and apical hook development in Arabidopsis seedlings (Hurný et al., 2020). The observed reduction of CK levels preceding the peak response of auxin content during waterlogging (Figure 2), indicates their interaction might be tissue-dependent and dynamically regulated in time during waterlogging (Figure 7).

### Waterlogging lowers levels of active CKs and their ribosides in leaves and petioles

CK are both synthesized locally and transported throughout the plant via the vascular tissue to systemically modulate plant development. To this end, riboside precursors of tZ type CKs are transported by the xylem, while precursors of iP and cZ type CKs are transported by the phloem (Hirose et al., 2008). These precursors are subsequently activated by LONELY GUY (LOG) to locally induce CK mediated processes (Kuroha et al., 2009). In addition, active CK such as tZ have also been found to be transported over longer distances to induce cytokinin responses at remote sites (Osugi et al., 2017). Given the significant reduction of especially these transportable riboside precursors, CK transport is likely rapidly downregulated during waterlogging (Figure 2 & Figure 7). As a consequence, and possibly together with a reduced CK biosynthesis, the pool of active CK is reduced in leaves of all ages. Furthermore, active CKs are conjugated through *N*-glucosylation (iP7G, cZ7G) and *O*-glucosylation (cZROG). On the other hand, cZR levels remain higher in older leaves, while DHZ7G formation stays limited. It has been shown before during senescence in Arabidopsis that CK *N*-glucosides, together with tZOG, increased in leaves (Šmehilová et al., 2016). Besides being inactive yet abundant storage forms, the above glucosides are emerging as points of CK control, possibly by regulating CK responses and homeostasis (Hallmark et al., 2020; Pokorná et al., 2021). Despite increasing evidence on transportability and possibly even reconversion into active CK (tZ7G) (Hošek et al., 2020), many questions on the exact function of these glucosides remain.

In tomato roots, the CK controlled *CYTOKININ OXIDASE/DEHYDROGENASE 2 (SlCKX2)* is strongly upregulated during waterlogging (De Ollas et al., 2021), indicating that CK degradation is activated in the roots. In contrast, waterlogging seems to repress the *CYP735A1/2* orthologs (Solyc02g094860, Solyc02g085880), involved in tZR and tZ biosynthesis, in tomato roots (De Ollas et al., 2021). A reduction in xylem CK levels and shootward transport could explain why these root-derived CK types are also drastically reduced in leaves during waterlogging (Figure 2 & 3). In common bean *(Phaseolus vulgaris)* and poplar *(Populus trichocarpa* x *P. deltoides)*, waterlogging has already been shown to reduce root-to-shoot CK fluxes (Neuman et al., 1990).

Despite multiple reports on the negative regulation of stress responses by CK, there are indications that their role in stress signaling might be more complex (O’Brien & Benková, 2013). Increased CK degradation by CKX and reduced CK signaling have been shown to enhance resilience in Arabidopsis to salt and drought stress (Abdelrahman et al., 2021; Nishiyama et al., 2011), while higher CK production through *pSAG12::IPT* expression enhances flooding resilience (Huynh et al., 2005; Zhang et al., 2000). Perhaps the actual stress response depends on dynamic changes of CK production and signaling. Huynh et al. (2005) reported a transition in shoot CK levels during waterlogging in Arabidopsis, consisting of an initial decline and followed by an accumulation (as observed for iP; Figure 2).

### Dampened CK homeostasis paving the way for epinastic bending in tomato?

Despite their role in cell proliferation, plant growth and stress resilience, CK have received surprisingly little attention for their potential in mediating leaf posture. Only recently, it has been proven that their degradation by OsCKX3 might actually regulate leaf angle in rice (Huang et al., 2022). In tomato, application of BA together with GA, reduces waterlogging-induced epinasty, indicating that either GA or CK (or both) are negative regulators of epinasty (Jackson & Campbell, 1979).

Given their close interaction during growth and development, CK and auxins might collectively control cell elongation to induce epinastic bending. It has been shown in Arabidopsis that belowground gravitropic bending of lateral roots is restrained by asymmetric CK signaling, indicating that CK and auxins can actually co-regulate organ bending (Waidmann et al., 2019) (Figure 7). In addition, CK induce the auxin/indole-3- acetic acid (Aux/IAA) gene *SHORT HYPOCOTYL2* (*IAA3/SHY2*) to optimize cell division and differentiation in roots (Dello Ioio et al., 2008). While shoot and root responses are structurally different, these examples provide a line of evidence for CK-mediated differential growth, by interfering with both cell elongation and cell-cycle in concert with auxins.

How could the observed CK reduction be related to waterlogging-induced epinasty? There is increasing evidence suggesting CKs promote ethylene biosynthesis through stabilization of ACC synthase (ACS) (reviewed by Hansen et al., 2009). However, in case of waterlogging, root-borne ACC is translocated to the shoot and conversion by ACO becomes rate-limiting (English et al., 1995; Jackson & Campbell, 1976). One of these ACOs *(ACO5)* is downregulated by CK treatment in Arabidopsis leaves (Brenner et al., 2005) and modulates the NAC transcription factor SPEEDY HYPONASTIC GROWTH (SHYG) to establish hyponastic growth during waterlogging (Rauf et al., 2013). Similarly, CK treatment of detached tomato leaves represses expression of *ACO5* (Solyc07g026650) in leaves of 35-day-old plants already after 2 h (Shi et al., 2013) (Figure 7). At least three members of the *ACO* family are upregulated during waterlogging in tomato (*ACO1*, *ACO3* & *ACO6*), while only one is downregulated (*ACO4*) (De Ollas et al., 2021). So far, the role of CK in the regulation of ethylene biosynthesis in tomato is unknown.

Further downstream, CKs potentially interact with DWARF (DWF) mediated BR signaling not only to regulate bud outgrowth in tomato through RESPONSE REGULATOR10 (RR10) (Xia et al., 2021). Leaves of DWF impaired mutants have a reduced ABA and ethylene biosynthesis and are hyponastic, while CKs are not unequivocally affected in both the defective and the DWF overexpressing mutant (Li et al., 2016).

### CK, ABA and sugar crosstalk as modulator of waterlogging-induced epinasty?

The contrasting dynamics of ABA and CK levels during waterlogging (Figure 2) could hypothetically transduce a signal to facilitate epinastic bending (Figure 7). A similar peak in ABA content, coinciding with a reduction of CK has been observed during waterlogging in Arabidopsis (Huynh et al., 2005). In addition, it has been shown that a decline of CK seems to sensitize Arabidopsis plants to ABA, while ABA inhibits CK biosynthesis, leading to complex feedback regulation (Nishiyama et al., 2011).

ABA and CK are closely intertwined with sugar signaling and sink-source balancing. This interface provides a possible point of crosstalk during waterlogging, as adequate carbohydrate allocation during low-oxygen stress is crucial for survival. Soluble sugar content increases drastically in shoots of waterlogged Arabidopsis plants (Huynh et al., 2005) and barrel clover (Lothier et al., 2020), indicating export to the roots is restricted and sugar homeostasis disturbed (Lothier et al., 2020). In turn, these changes in carbohydrate levels could induce hormone-driven processes. Both CK and ABA for example interact with SNF Related Kinase (SnRK) – TARGET OF RAPAMYCIN (TOR) mediated sugar signaling to control growth (Dong et al., 2015; Wang et al., 2018) and antagonistically regulate stress responses (Huang et al., 2018). This sugar-hormone crosstalk provides another hypothetical framework for waterlogging-induced and ontogenically-regulated epinasty that needs further investigation (Figure 7).

