## Supplementary Material for "Waterlogging shifts ontogenic hormone dynamics in tomato leaves and petioles"

### Supplemental material

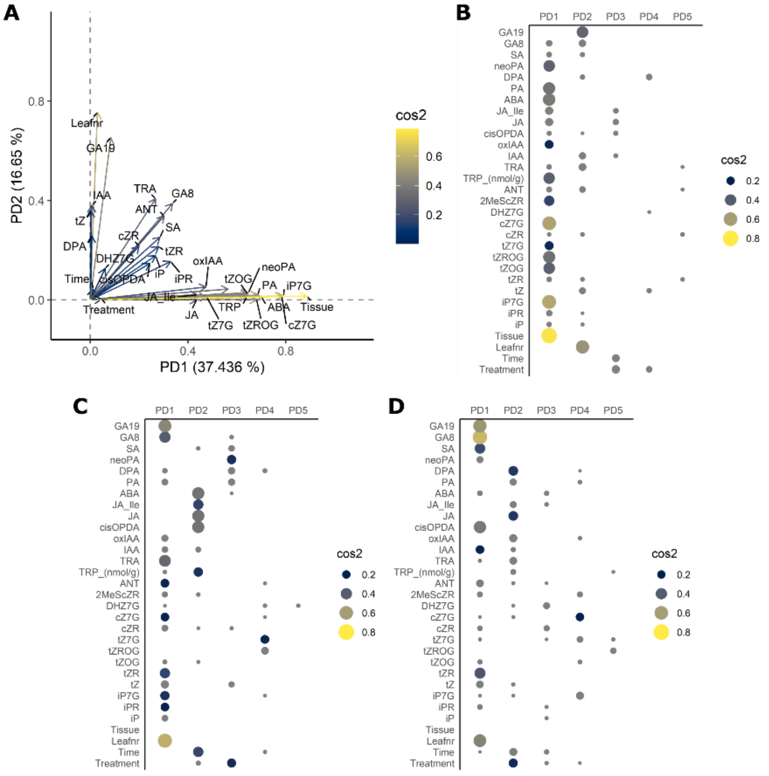


Supplemental Figure S1: Factor analysis of mixed data (FAMD) analysis of normalized hormone levels of tomato leaves and petioles during waterlogging. (A) Variable plot, including design variables and hormone variables. (B – D) Quality of representation of the variables on the FAMD principal dimensions (PD) in (B) leaves and petioles together, and (C) leaves and (D) petioles separately.
